# The altered T cell landscape in Systemic Sclerosis patients is characterized by dysfunctional type 1 immunity

**DOI:** 10.64898/2025.12.10.693457

**Authors:** Victoria Volfson-Sedletsky, Hannah A. DeBerg, Mitch L. Fahning, Ian R. Rifkin, Andreea M. Bujor, Daniel J. Campbell, Peter A. Morawski, Anna C. Belkina, Hans Dooms

**Affiliations:** Arthritis and Autoimmune Diseases Center, Rheumatology Section, Department of Medicine, Boston University Chobanian and Avedisian School of Medicine, Boston, MA; Department of Virology, Immunology & Microbiology, Boston University School of Medicine, Boston MA; Center for Systems Immunology, Benaroya Research Institute, Seattle, WA; Center for Fundamental Immunology, Benaroya Research Institute, Seattle, WA; Renal Section, Department of Medicine, Boston Medical Center, Boston, MA; Renal Section, Department of Medicine, VA Boston Healthcare System, Boston, MA; Department of Immunology, University of Washington School of Medicine, Seattle, WA; Department of Pathology, Boston University School of Medicine, Boston, MA; Department of Immunology and Genomic Medicine, National Jewish Health, Denver, CO; Department of Immunology and Microbiology, University of Colorado Anschutz Medical Campus, Aurora, CO

## Abstract

**Background:** T cells in patients with chronic autoimmunity show features of dysfunction and exhaustion. However, the functionally-defined T cell subsets affected by these changes remain poorly characterized. Here, we sought to reveal aberrations in the composition, phenotype and function of canonical T cell subsets in the blood of Systemic Sclerosis (SSc) patients, compared to healthy subjects and Systemic Lupus Erythematosus (SLE) patients.

**Methods:** We developed a novel multidimensional flow-cytometry panel to simultaneously detect lineage-defining transcription factors, co-inhibitory receptors and other functional markers to characterize T cell subsets without *in vitro* restimulation. We compared T cell landscapes in SSc and SLE patients, and healthy subjects, using Optimized t-SNE and PhenoGraph algorithms. Cytokine production in patient samples was determined using intracellular cytokine staining. Transcriptomic analysis was performed to assess the role of IFN-γ in fibroblasts.

**Results:** The data show altered distribution and aberrant functional states of transcription factor-defined T cell subsets in patients with autoimmunity, with shared and unique features between diseases and their subtypes. Strikingly, SSc and SLE patients showed a severe deficiency in subsets of CD8^+^ and CD4^-^CD8^-^ T cells that expressed T-bet, which is critical for IFN-γ-dependent type 1 immunity, and exhibited features of exhaustion. Moreover, TIGIT^+^Foxp3^+^ regulatory T cells, known to suppress Th1 responses, were selectively increased in SSc. Functionally, T cells from SSc patients produced less IFN-γ, a cytokine that suppresses pro-fibrotic gene expression in fibroblasts.

**Conclusion:** Our study demonstrates that SSc patients have a defective IFN-γ-producing T cell compartment that may lead to reduced anti-fibrotic T cell activity, enabling chronic fibrosis development.

**Funding:** National Institute of Allergy and Infectious Diseases/NIH, National Institute of Arthritis and Musculoskeletal Diseases/NIH, National Heart, Lung and Blood Institute/NIH, National Scleroderma Foundation, National Jewish Health

## Introduction

Systemic Sclerosis (SSc) or Scleroderma is a complex autoimmune disease characterized by immune dysregulation, vasculopathy, and progressive fibrosis of the skin and internal organs. While the precise role of T cells in SSc pathogenesis remains incompletely understood, a growing body of evidence implicates their involvement in promoting disease pathology (1, 2). Multiple T cell subsets, including Th1, Th2, Th17, Tregs, Tfh, cytotoxic CD4^+^ T cells, and CD8^+^ T cells have been implicated in the disease process (3–9). The role of these T cell subsets in fibrosis may be attributable to their capacity for pro- and anti-fibrotic cytokine secretion, cytotoxic activities, immune regulation and autoantibody production. Many of these functions are regulated by co-stimulatory (co-SR) and co-inhibitory (co-IR) receptors. We and others have shown that some co-IRs are increased on CD4^+^ and CD8^+^ T cell subsets in SSc (10), as well as in many other autoimmune diseases (11–14). However, a detailed analysis of the functionally-defined T cell lineages and their subsets that are affected by specific patterns of co-IR and co-SR expression has not been performed. One challenge to link co-SR and co-IR expression to functional subsets is the need to restimulate T cells in *vitro* to define their cytokine production profiles. This process modulates co-SR/-IR expression, making assignment of the phenotypes to the original *ex vivo* subsets potentially inaccurate. To address this problem, we designed a novel multidimensional spectral flow cytometry panel that combines staining for transcription factors defining cytokine-competent T cell lineages with various surface and intracellular markers indicative of T cell fitness and functional state. This approach allowed to directly quantify the presence of functionally-defined CD4^+^, CD8^+^ and CD4^-^CD8^-^ T cell subsets in peripheral blood of patients with diffuse cutaneous SSc (dcSSc), limited cutaneous SSc (lcSSc), and SLE, and define their phenotype. Our study revealed significant differences in the T cell landscape of SSc patients, SLE patients and healthy subjects, mainly characterized by a striking defect of specific T-bet-expressing T cell populations in SSc and SLE. Functionally, memory T cell subsets from SSc patients showed reduced IFN-γ production after *in vitro* restimulation. This deficiency in frequency and function of T cells that provide type 1 immunity may significantly impact SSc disease course, since we show that IFN-γ reduces collagen production in fibroblasts. We also found an increase in TIGIT-expressing Tregs that may further suppress IFN-γ-producing T cells (15). Hence, our findings suggest that a defective IFN-γ compartment leads to weakened anti-fibrotic activities in SSc patients.

## Results

### Significant phenotypic heterogeneity within T cell lineages defined by transcription factor expression in human peripheral blood

We designed a multidimensional flow cytometry panel comprising of 20 markers to perform a comprehensive analysis of T cells expressing lineage-defining transcription factors in combination with markers of activation and dysfunction in T cells, including T-bet, GATA-3, RORγt, Bcl-6, Foxp3, TOX, PD-1, TIGIT and CD226 (Suppl. Table 1A). All antibody clones and concentrations were extensively tested and optimized using methods we previously published (16). To validate that the intracellular markers faithfully identified distinct T cell subsets, we utilized opt-SNE analysis (17), generating heatmaps illustrating the expression levels of these markers (Fig. 1A, B, C). Canonical lineage-defining transcription factors showed expected expression patterns within CD4^+^, CD8^+^ and CD4^-^CD8^-^ T cell populations: for example, T-bet, GATA3, and RORγt were present in all three T cell populations, while Foxp3 expression was confined to CD4^+^ and CD4^-^CD8^-^ T cells. Among CD4^+^ T cells, the majority of transcription factors were co-expressed with the CD45RO memory marker, confirming they represented subsets of antigen-experienced T cells. In CD8^+^ and CD4^-^CD8^-^ T cells some transcription factor-expressing subsets were detected within CD45RO^-^ T cells. Although CD45RA was not included in our panel, these CD45RO^-^ subsets are likely part of the naïve T cell population or may be terminally differentiated effector memory T cells that re-express CD45RA (TEMRAs)(18). Co-expression of most lineage-defining transcription factors was minimal, confirming the identification of discrete subsets (Fig. 1A, B, C). Thus, we have developed a unique flow cytometry panel that combines staining for an extensive number of intracellular proteins with viability, proliferation, and functional markers, thus allowing to faithfully and simultaneously identify all major functional T cell subsets in human blood and assess their biological state.

**Figure 1:**
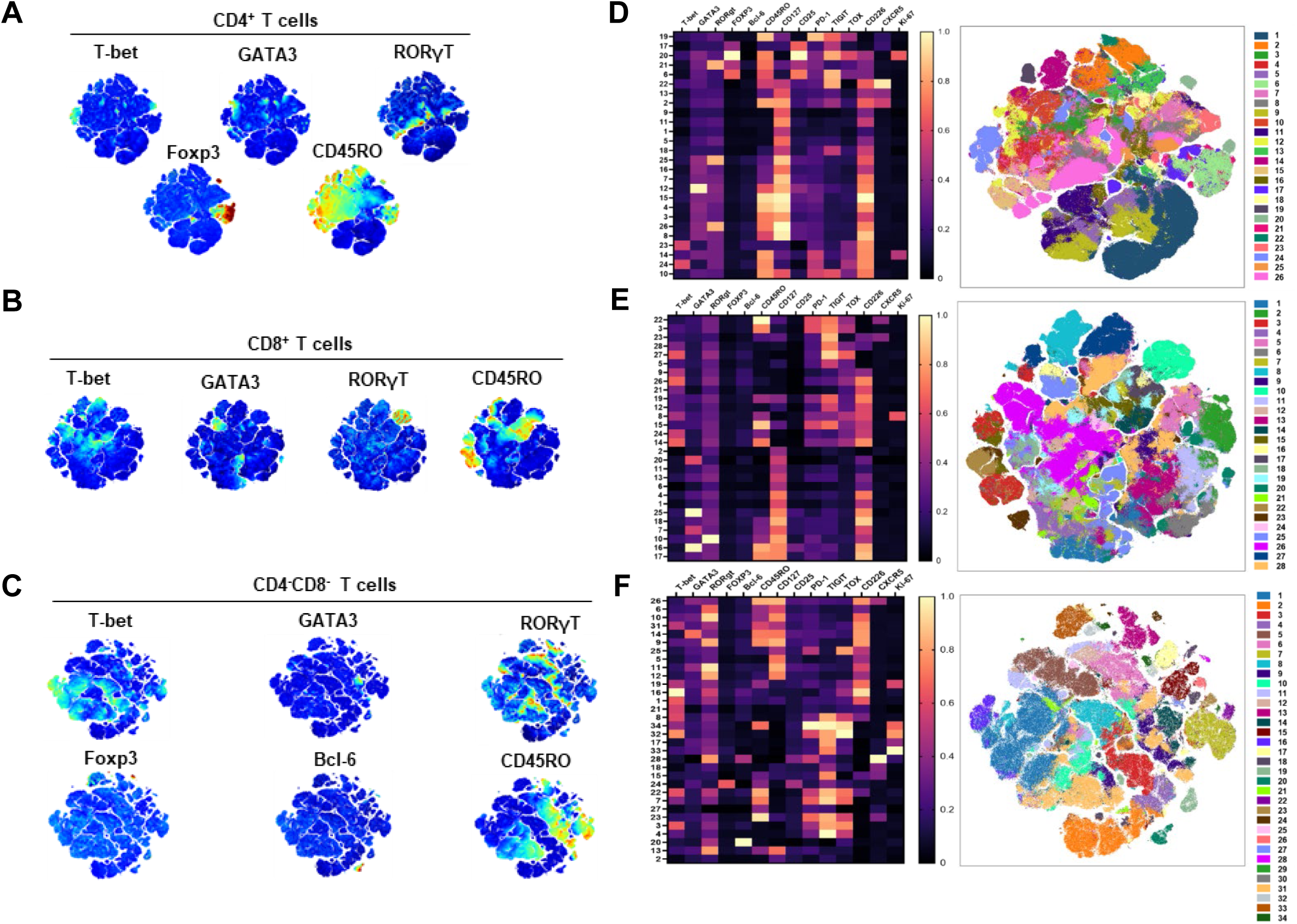
Identification of T cell subsets in human peripheral blood based on lineage-defining transcription factors reveals distinct functional phenotypes within canonical subsets. Peripheral blood samples of healthy controls (HC), dcSSc, lcSSc and SLE subjects were analyzed with a flow cytometry panel for full spectrum flow cytometry (FSFC) (Suppl. Table 1). Data were analyzed using opt-SNE algorithm and PhenoGraph clustering. Transcription factor expression patterns are shown as heatmaps overlaid on opt-SNE plots of live single CD3+ and (A) CD4+ T cells, (B) CD8+ T cell and (C) CD4^-^CD8^-^ T cells. Normalized expression levels of intracellular and surface proteins in PhenoGraph clusters are shown in heatmaps for (D) CD4+ T cells, (E) CD8+ T cell and (F) CD4^-^CD8^-^ T cells. Numbers represent the different, hierarchically arranged clusters. Values for every marker are normalized to minimum and maximum values in each column across CD4+, CD8+ and CD4^-^CD8^-^ T cell data for comparable analysis. Data includes all samples concatenated.

To generate a detailed map of the phenotypical and functional diversity within T cell subsets, we employed the PhenoGraph algorithm that clusters cell populations in an unbiased manner based on their expression profiles of all the markers of interest.

PhenoGraph clustering identified 26 distinct clusters in CD4^+^, 28 clusters in CD8^+^, and 34 clusters in CD4^-^CD8^-^ T cells from all concatenated human peripheral blood samples (Fig.1D, E, F). To identify the markers that defined each cluster, Median Fluorescence Intensities (MFI) within the PhenoGraph-generated clusters of CD4^+^, CD8^+^ and CD4^-^CD8^-^ T cells were plotted on a hierarchically arranged heatmap (Fig 1D, E, F). Each marker expression was normalized across CD4^+^, CD8^+^ and CD4^-^CD8^-^ T cell populations to be able to compare differences in expression between these three T cell subsets. The combination of transcription factors and functional surface receptors detected with our panel revealed previously unknown phenotypes within canonical T cell subsets in human peripheral blood, allowing to compare the frequencies of these subsets in healthy and diseased individuals. Some noted features included that CD8^+^ and CD4^-^CD8^-^ T cells contained more individual clusters defined by T-bet expression than CD4^+^ T cells, while, as expected, Foxp3^+^ T cells were largely confined to CD4^+^ T cells. Interestingly, large areas within the CD45RO^+^ T cell region did not show expression of a canonical transcription factor, suggesting the prevalence of uncommitted, reverted or undefined cell populations. Many clusters expressed the co-SR CD226, while its competing co-IR TIGIT, which binds to the same ligands, showed more limited cluster distribution. GATA-3 was only highly expressed in one cluster in CD4^+^ and CD4^-^CD8^-^ T cells but multiple among CD8^+^ T cells. CD4^-^CD8^-^ T cells contained several clusters positive for RORγt, while CD4^+^ and CD8^+^ T cells showed fewer diversity in RORγt^+^ cells. This unbiased clustering analysis hence documented the T cell landscape in human peripheral blood and revealed previously unknown details of T cell subset composition.

### Features of a compromised type 1 T cell compartment dominate the altered T cell landscape in SSc and SLE patients

Next, we leveraged our data set to assess differences in cluster frequencies between healthy subjects, SSc patients and SLE patients. For this analysis, we used a carefully characterized patient cohort representing SSc disease heterogeneity. SSc and SLE cohort characteristics are shown in Table 1, demonstrating that healthy and SSc cohorts were largely age- and sex-matched, while SLE patients tended to be younger, consistent with typically earlier disease onset. The SSc population consisted of lcSSc (n=21) and dcSSc (n=22) patients with and without SSc Interstitial Lung Disease (SSc-ILD) and Pulmonary Arterial Hypertension (PAH). It is also important to note that patients that were not taking immunosuppressive drugs and patients that were treated with these drugs were both represented, allowing to evaluate the impact of drug treatment on the T cell landscape. First, we assured that the different cohorts in our data (healthy subjects, dcSSc, lcSSc and SLE) showed comparable overall cell distribution patterns. Major differences in cell distribution and cell population ratios between the groups may indicate technical issues and can result in an inaccurate analysis, both biological and statistical. To exclude this, we generated opt-SNE maps that were overlayed with cell densities for each cohort for all three T cell lineages (Suppl. Fig. 1). Each cohort contained an equal number of CD4^+^ (1 x 10^6^ cells), CD8^+^ (1 x 10^6^ cells) and CD4^-^CD8^-^ (250,000) T cells. We did not observe large regions that were only present in some cohorts but not others, indicating the stained samples were intact and the staining quality comparable, allowing us to execute analysis of all four cohorts. Interestingly, a shift was observed towards increased presence of CD45RO^+^ clusters in SSc and SLE patients, which indicates a more “aged” and/or active immune system as has been described for systemic autoimmune diseases (19).

**Table 1:**
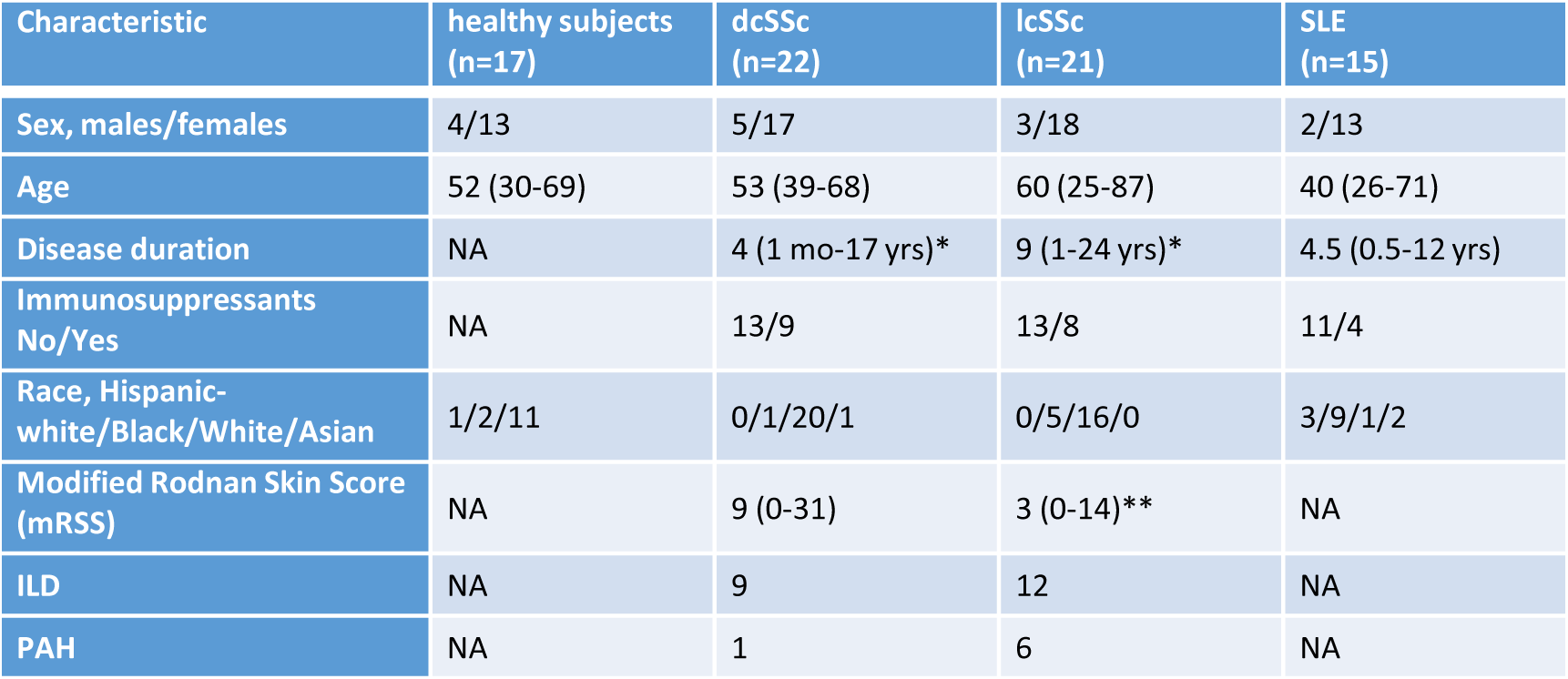
Clinical information of healthy subjects, SSc, and SLE patients. * No MRSS reported for 1 dcSSc patient and 3 lcSSc patients ** disease duration is not available for 3 lcSSc patients

Next, we determined the frequency of each cluster identified by the automated clustering algorithm (Fig. 1) in healthy subjects, SSc, and SLE patients using OMIQ software. First, to evaluate the paradigm that fibrosis in SSc is driven by type 2 immune cells (20, 21), we analyzed T cell subsets expressing the canonical type 2 transcription factor GATA-3. Surprisingly, we found that within CD4^+^ T cells, the frequencies of the CD45RO^+^ GATA-3-expressing clusters 12 (GATA3^high^) and 15 (GATA3^low^) were not significantly different in SSc patients vs healthy subjects (Fig. 2A). In SLE patients, GATA-3-expressing cells were significantly reduced compared to both healthy subjects and dcSSc patients. Similarly, within CD8^+^ T cells, GATA-3-expressing clusters were not altered in SSc patients, but reduced in SLE patients (Fig. 2B). The main GATA3^+^ subset within CD4^-^CD8^-^ T cells showed no differences between the cohorts (data not shown). Hence, SSc patients did not show an increased bias towards a type 2 immune landscape in peripheral blood, as concluded from a lack of expansion of GATA-3^+^ subsets. This appears to contrast with reports that show an increase in Th2/Tc2 cytokines in SSc patients compared to healthy subjects (22–24). Hence, it is feasible that GATA-3^+^ T cells are not more numerous in SSc but exhibit a lower threshold for cytokine production.

**Figure 2:**
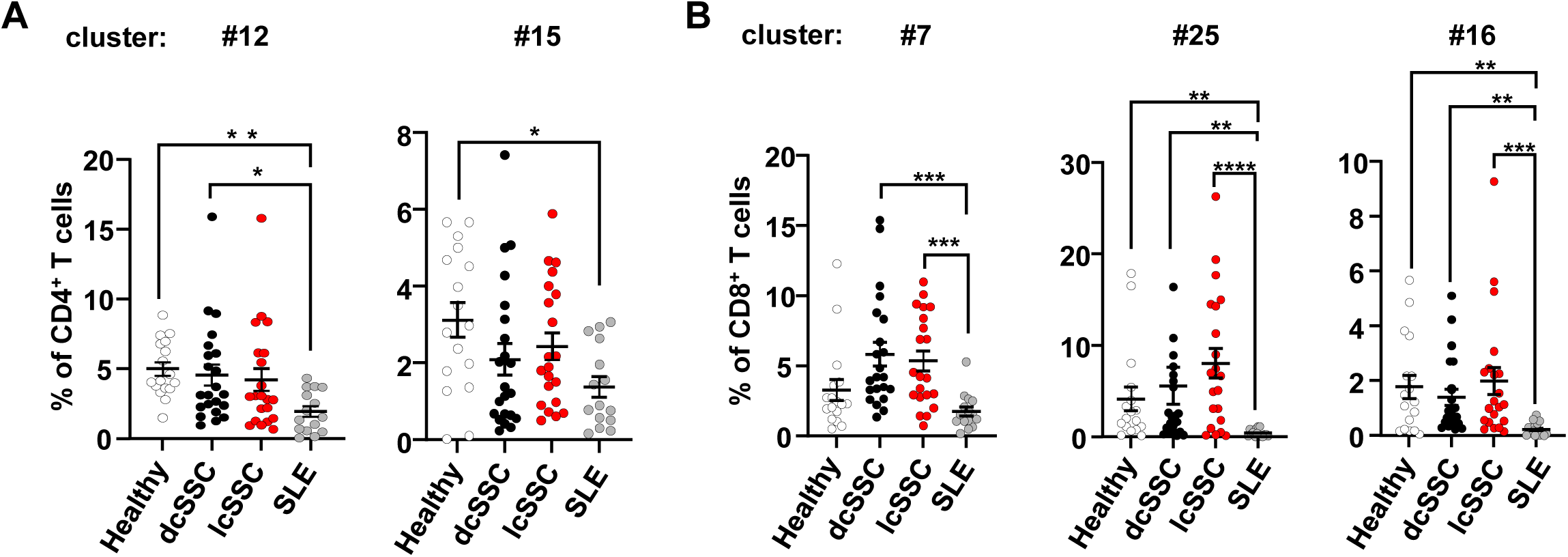
SSc patients do not shown major changes in GATA-3-expressing T cells. Frequency of GATA-3-expressing clusters was determined using markers that characterized the cluster as identified in heatmaps from Fig.1D, E, using OMIQ software. Cell frequencies of indicated clusters within parental populations of (A) CD4+, and (B) CD8+ T cells, in healthy controls (HC) (n=17), dcSSc (n=23), lcSSc (n=21), SLE (n=15) patients are shown. Bar graphs show percentages of gated cells of indicated clusters with mean and SEM. Each symbol represents a blood sample from an individual donor. Data was statistically analyzed with the Kruskall-Wallis test followed by Dunn’s multiple comparisons test. *p<0.05, **p<0.01, ***p<0.001, ****p<0.0001.

To assess prevalence of type 1 immunity, we analyzed T-bet-expressing clusters. Within CD4^+^ T cells, the two main T-bet^+^ clusters, 23 and 24, which were distinguished by expression of CD45RO and represented each about 1% of CD4+ T cells, were increased in SLE patients but not in SSc patients, compared to healthy subjects (Fig. 3A). T-bet directs Th1 lineage commitment and T-bet-expressing CD4^+^ T cells are known to be increased in SLE patients and may have a pathophysiological role in the disease, underscoring our findings confirm previous studies (25). Within CD8^+^ T cells, many T-bet-expressing clusters were identified, with 14, 26, and 27 representing the most cells. The CD45RO^+^ T-bet^+^ cluster 14 that expressed CD226, TIGIT and TOX, and the CD45RO^-^ T-bet^+^ cluster 26, characterized by CD226, TOX and PD-1 expression, showed no difference in SSc patients and a minor increase in SLE patients, not reaching statistical significance (data not shown). Strikingly, cluster 27, composed of T-bet^+^ CD45RO^-^ CD127^-^CD226^-^TOX^hi^TIGIT^hi^ CD8^+^ T cells, was drastically reduced in SSc and SLE patients compared to healthy subjects (Fig. 3B). This cluster represented the largest number of T-bet-expressing CD8+ T cells in healthy subjects, and the high levels of TOX and TIGIT, along with the absence of co-stimulatory molecule CD226, suggests this is an exhausted population of Tc1 cells. Similarly, in CD4^-^CD8^-^ T cells, cluster 16, which expressed high levels of T-bet, was more than 10-fold reduced in all three disease types (dcSSc, lcSSc and SLE) compared to healthy subjects (Fig. 3C).

**Figure 3:**
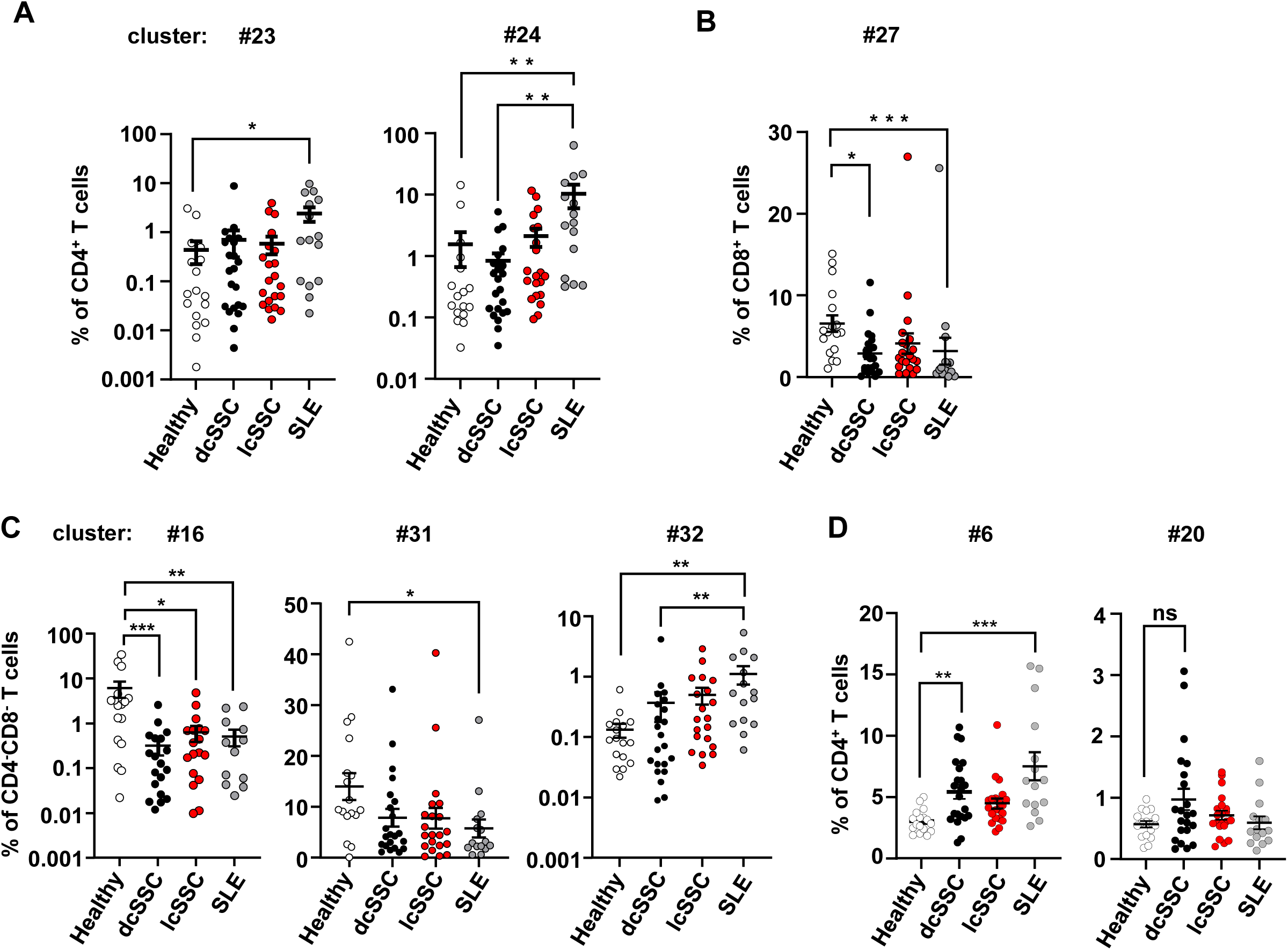
SSc and SLE patients exhibit features of reduced type 1 immunity. Frequency of T-bet- and Foxp3-expressing clusters was determined using markers that characterized the cluster as identified in heatmaps from Fig.1D, E, F, using OMIQ software. Cell frequencies of indicated T-bet-expressing clusters within parental populations of (A) CD4+, (B) CD8+, and (C) CD4^-^CD8^-^ T cells, and cell frequencies of indicated Foxp3-expressing clusters within CD4+ T cells (D) are shown for healthy controls (HC) (n=17), and dcSSc (n=23), lcSSc (n=21), SLE (n=15) patients. Graphs show percentages of gated cells of indicated clusters with mean and SEM. Each symbol represents a blood sample from an individual donor. Data was statistically analyzed with the Kruskall-Wallis test followed by Dunn’s multiple comparisons test. *p<0.05, **p<0.01, ***p<0.001, ****p<0.0001

Cluster 31 was also reduced in SLE and showed a similar trend in SSc. Staining with anti-Vδ2 TCR antibodies revealed that these clusters belonged to the Vδ2^+^ subset of γδ T cells, the most ubiquitous γδ T cell population in human blood (26). Cluster 32 on the other hand, representing Vδ1^+^ γδ T cells, was increased in SLE and exhibited a highly activated phenotype, characterized by co-expression of PD-1, TIGIT, TOX and the proliferation marker Ki-67 (Fig. 3C). Hence, we conclude that SSc and SLE patients show features of an altered type 1 immune compartment that is not characterized by a broad and indiscriminate reduction of T-bet-expressing T cells but by specific deficiencies in defined subsets. Moreover, further support for an altered type 1 immune response in SSc and SLE patients came from the finding that frequencies of Foxp3+ Tregs expressing high levels of TIGIT (cluster 6), a subset that has been associated with preferential suppression of Th1 but not Th2 cells, were increased (Fig. 3D) (15, 27). A second TIGIT+ Treg cluster, #20, also showed a trend to be increased in dcSSc, but not lcSSc and SLE patients. The main difference between these 2 clusters is that #20 expressed Ki67 and TOX, indicating a highly activated, proliferating population of TIGIT^+^ Tregs (Fig. 3D). Treg clusters (17 and 21) that expressed low levels of TIGIT did not show a difference between healthy subjects and patients (data not shown). In conclusion, analysis of T cell subsets expressing GATA-3, T-bet, or Foxp3 in human peripheral blood revealed that changes are not uniform across all cells expressing these lineage-defining transcription factors. Instead, specific, previously undefined subsets within these populations are altered in frequencies in a disease-specific manner, and these changes are indicative of a dysregulated type 1 immune response in SSc and SLE.

### Select type 17 T cell subsets are sensitive to immunosuppressive drugs in SSc patients

Next, we analyzed the composition of RORγt-expressing CD4^+^, CD8^+^ and CD4^-^CD8^-^ T cells in our patient cohorts. In CD4^+^ T cells, RORγt expression defines the Th17 lineage (28). Th17 cells are expanded in peripheral blood of SSc patients but their role in the disease remains poorly defined (7, 29–31); in SLE, Th17 cells are thought to exacerbate the disease (32, 33). In our study, we found that three CD4^+^ clusters expressed high levels of RORγt: cluster 21 co-expressed Foxp3 and has been discussed above, clusters 25 and 26 were distinguished by CD45RO expression. CD45RO^-^ RORγt-expressing T cells constituted on average <2% of CD4^+^ T cells and were increased in dcSSc and SLE patients compared to lcSSc patients, which had very low levels of these cells (Fig. 4A). Interestingly, separating SSc patients based on immunosuppressive drug treatment revealed that the higher frequencies of RORγt^+^ T cells in dcSSc were driven by patients that were not taking immunosuppressive drugs (Fig. 4A). Treatment with immunosuppressants reduced RORγt^+^ T cells back to levels seen in healthy subjects and lcSSc patients, showing specific expansion of this population in dcSSc patients is sensitive to immunomodulation. Cluster 26, representing CD45RO^+^ effector/memory RORγt^+^ T cells showed only minor differences between patient cohorts (data not shown).

**Figure 4:**
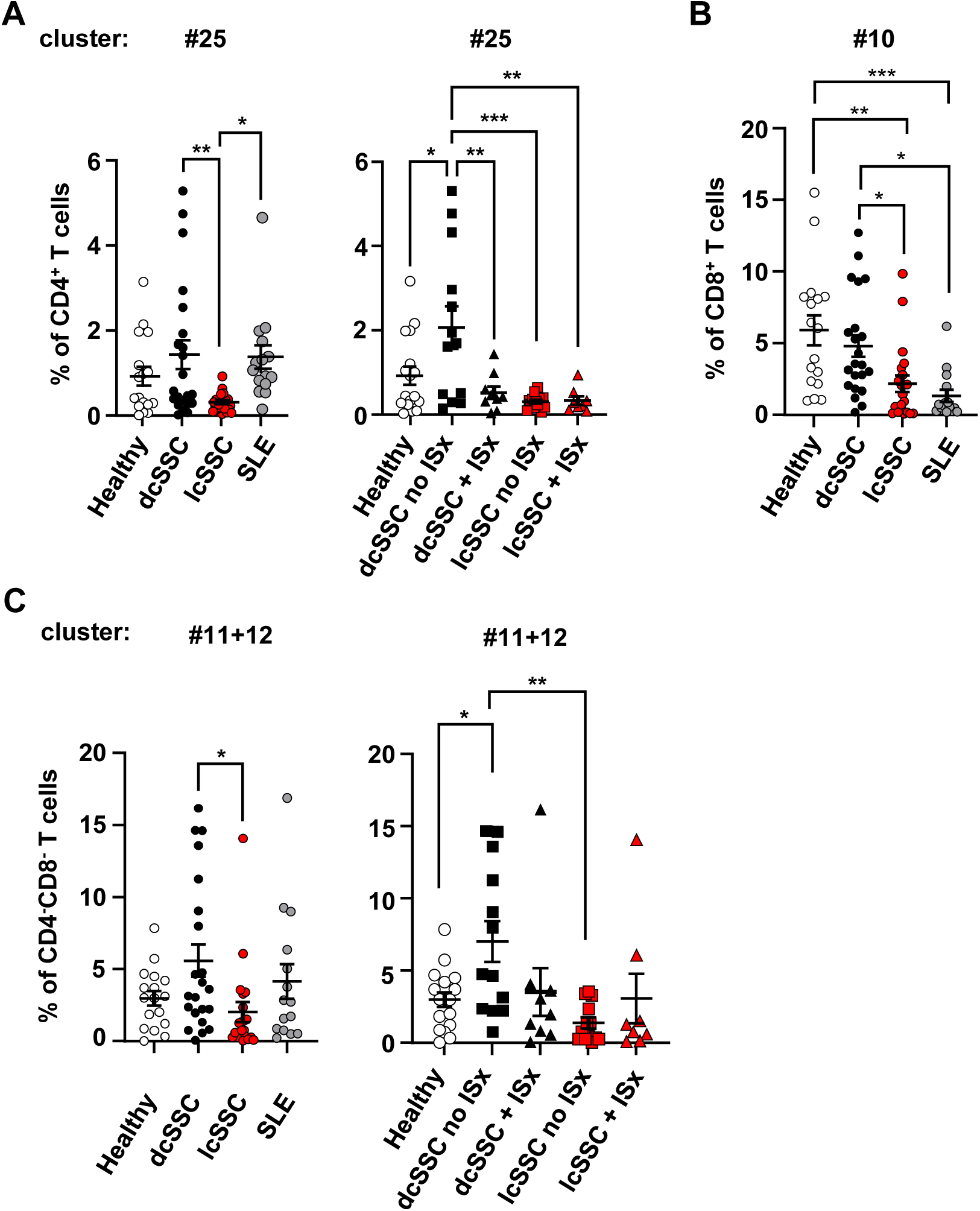
RORγt-expressing T cell clusters show disease-specific changes in frequency and are sensitive to immunomodulation in dcSSc patients. Frequency of RORγt-expressing clusters was determined using markers that characterized the cluster as identified in heatmaps from Fig.1D, E, F, using OMIQ software. Cell frequencies of indicated RORγT-expressing clusters within parental populations of (A) CD4+, (B) CD8+, and (C) CD4^-^CD8^-^ T cells, in healthy controls (HC) (n=17), dcSSc (n=23), lcSSc (n=21), SLE (n=15) patients are shown. In the right panels of (A) and (B) data from dcSSc and lcSSc patients separated based on immunosuppressant treatment (ISx) is shown. Bar graphs show percentages of gated cells of indicated clusters with mean and SEM. Each symbol represents a blood sample from an individual donor. Data was statistically analyzed with one-way ANOVA followed by Tukey’s multiple comparisons test. *p<0.05, **p<0.01, ***p<0.001, ****p<0.0001

In CD8^+^ T cells, RORγt expression defines Tc17 cells, which are a relatively recently discovered population of cells that has been shown to be involved in autoimmune, infection and cancer settings (34). Although the proportion of Tc17 cells was reported to be increased in blood from SLE patients (32), our study is, to our knowledge, the first to identify and quantify Tc17 cells in SSc patients. We observed a significant reduction in Tc17 cells in lcSSc patients, compared to healthy subjects and dcSSc patients (cluster 10) (Fig.4B). Surprisingly, in contrast to previous reports in the literature, we observed a reduction of Tc17 cells in SLE patients as well.

Within the CD4^-^CD8^-^ T cell population many RORγt^+^ clusters were detected, which varied in CD45RO, CD127 and CD226 expression (data not shown). However, most of these clusters did not differ in frequency between the cohorts. Clusters 11 and 12 were combined due to their similarity in RORγt^high^ and GATA3^low^ expression and were significantly increased in dcSSc patients compared to lcSSc (Fig. 4C). This increase was sensitive to immunosuppressant treatment (Fig. 4C). Notably, these clusters did not co-express any of the γδ T cell markers, hence presumably belong to a subset of iNKT cells that is designated as NKT17 (35). Interestingly, sensitivity to immunosuppressant drugs appeared to be mainly a feature of RORγt-expressing T cell subsets as analysis of the other transcription factor-expressing T cell subsets did not show a significant difference with and without treatment.

### Additional aberrant features of the T cell landscape in SSc and SLE

Several additional observations of interest were made, including a significant reduction of a cluster (#2) of CXCR5-expressing CD45RO^+^ CD4^+^ T cells in dcSSc an SLE, while lcSSc patients had normal levels of these cells, indicating a possible disease-specific role in dcSSc and SLE (Suppl. Fig. 2A). This cluster likely represents a population of blood memory T follicular helper (Tfh) cells, which often lack Bcl-6 expression (36).

Elevated levels of Tfh cells have been found in SSc patients and correlated with lung disease severity (37, 38). Within the CD4^-^CD8^-^ T cell population, a subset that expressed Bcl-6 in combination with a detectable level of RORγt (cluster 20) was significantly increased in SLE patients, and perhaps marginally in lcSSc as well (Suppl. Fig. 2B). This subset did not express Vδ1 or Vδ2 suggesting that these are likely iNKT cells. Although the existence of follicular helper iNKT cells that express Bcl-6 (39, 40), as well as RORγt^+^ iNKT cells (41, 42), is well established, there are no available reports of iNKT subsets that co-expresses both Bcl-6 and RORγt. Finally, we analyzed differences in clusters identified in Figs. 1D, E, and F that were not defined by transcription factor expression but showed features of interest, including sensitivity of some clusters to immunosuppressant treatment; notable findings are shown in Suppl. Fig. 3.

### Reduced production of the anti-fibrotic cytokine IFN-γ by circulating T cells from *SSc patients*

Our finding that SSc patients showed irregularities in T-bet-expressing T cell subsets and Tregs, prompted the question whether this was associated with deficiencies in production of the signature type 1 cytokine IFN-γ. Reduced IFN-γ production may have a profound impact on the control of fibrosis as it has been reported this cytokine has anti-fibrotic properties (43–47). To assess IFN-γ production in SSc donor T cells, we used blood samples curated from a cohort of SSc subjects at the Benaroya Research Institute who were not treated with immune-modifying drugs (48). Clinical characteristics of this cohort have been previously published (48). Pan-T cells from these SSc patients (n=17) and sex- and age-matched healthy controls (n=15) were isolated and stimulated with PMA/ionomycin followed by intracellular cytokine staining. We observed no major differences in total T cell or memory population frequencies between study groups (Suppl. Fig. 4A-D). Consistent with a defect in type 1 immunity, we found that memory CD4^+^ and CD8^+^ T cells from SSc subjects showed a marked reduction in IFN-γ production (Fig. 5A, 5B). We also observed a small but significant increase in IL-13 production by memory CD8^+^ T cells, but not CD4^+^ T cells (Fig. 5A, 5C), consistent with previous studies (1, 9).

**Figure 5:**
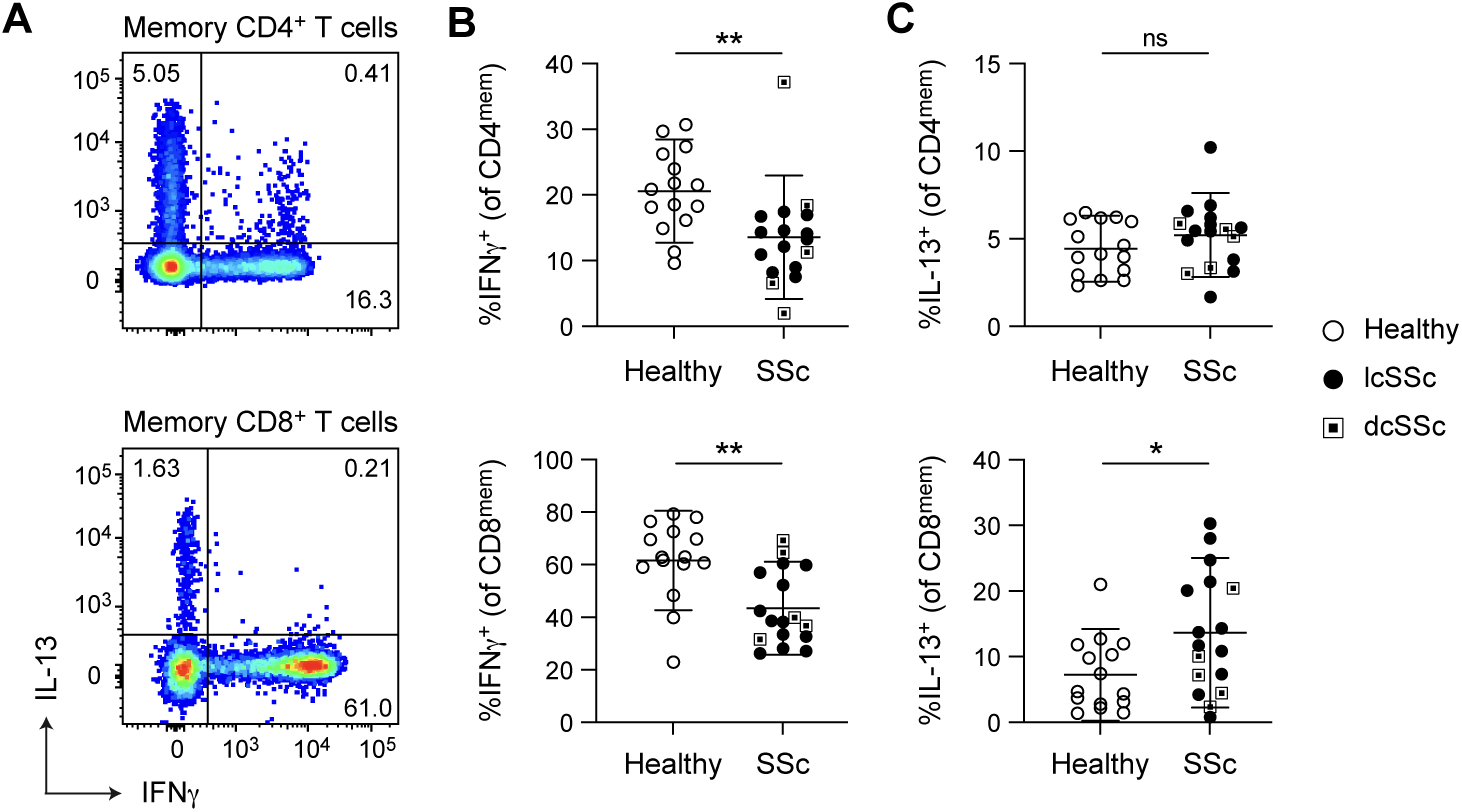
Peripheral blood T cells from SSc patients show reduced production of the anti-fibrotic cytokine IFN-γ. (A-C) Intracellular cytokine analysis of memory CD4+ (top) and CD8+ (bottom) T cells from healthy controls (n=15, open circles) and SSc (n=17, lcSSc: black circles; dcSSc: squares) donors following PMA/ionomycin stimulation. (A) Representative plots of indicated memory T cells showing IFN-γ and IL-13 expression. (B-C) Frequencies of cytokine-producing memory T cells across subject groups showing (B) IFN-γ+ and (C) IL-13+ cells as a percentage of CD4+ or CD8+ memory T cell subsets. Each point represents an independent biological replicate. Statistical significance: *p < 0.05, **p < 0.01, ns = not significant (unpaired t test or Mann-Whitney U test based on Shapiro-Wilk distribution testing). Error bars show mean ± SD.

To determine if reduced IFN-γ production by T cells could translate into diminished anti-fibrotic activity, consistent with the cytokine’s reported role in connective tissue disorders (44, 45), we compared gene expression in healthy human fibroblasts exposed to recombinant IFN-γ, IL-13, or IL-17a for 24 hours. Dimensionality reduction and differential gene expression analyses revealed that IFN-γ had the greatest impact on fibroblast gene expression, with more subtle effects induced by IL-13 and IL-17a (Fig. 6A, Suppl. Fig 5A-5C). We next assessed public transcriptomic signatures shared across fibrotic diseases including SSc (49, 50) and found a significant overlap between these and our cytokine-dependent effects on dermal fibroblasts (Fig. 6B). IFN-γ promoted a strongly expected signature of ISGs and gene pathways, while inhibiting fibrosis-associated gene pathways that were induced by IL-13, such as collagen formation and extracellular matrix organization (Fig. 6B-6C, Suppl. Fig. 5A-5B). Using a curated list of common ISG and fibrosis-associated genes we found further evidence for an opposing relationship between IFN-γ and IL-13 effects in fibroblasts (Fig. 6D-6E).

**Figure 6:**
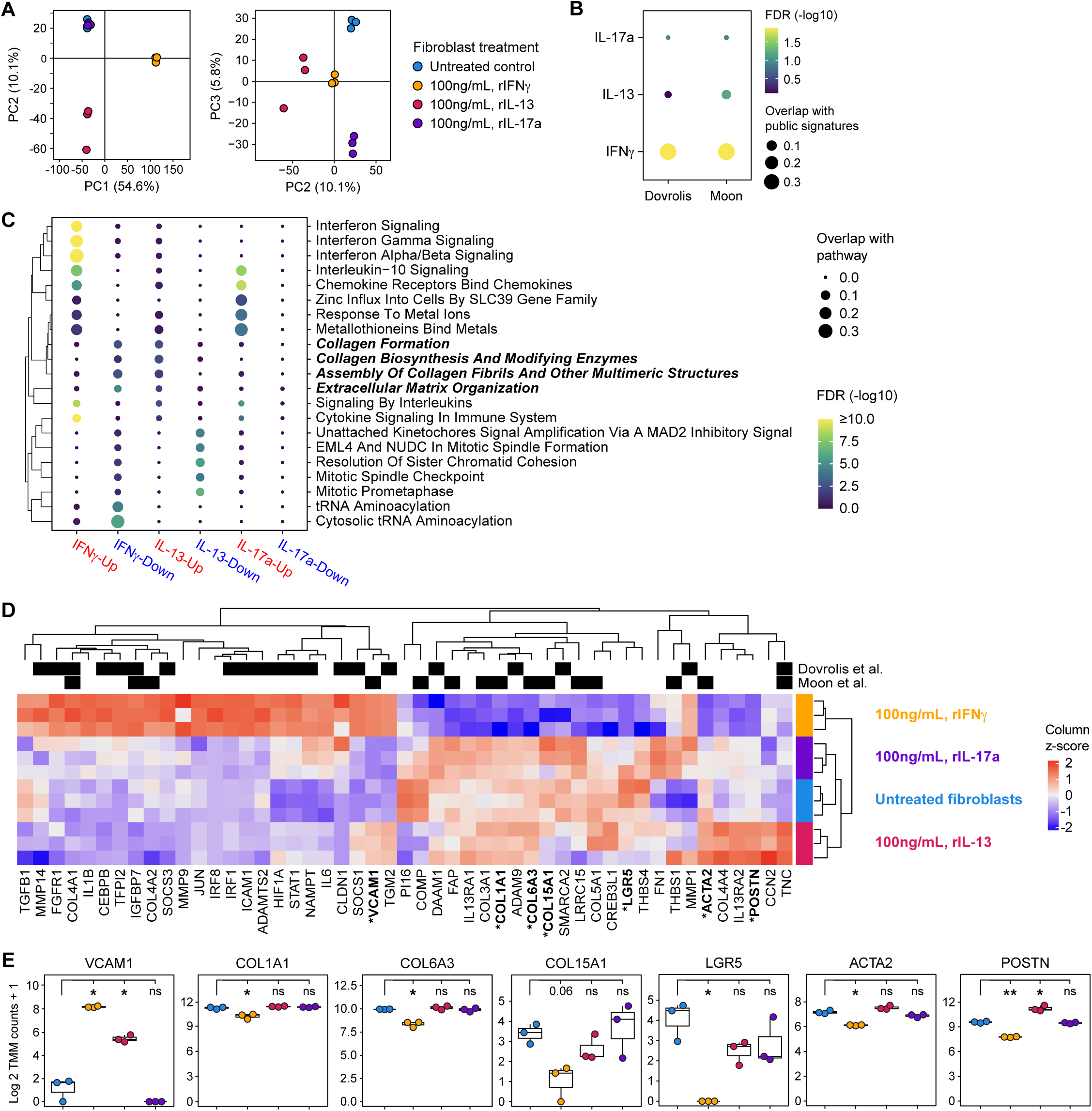
Cytokine-mediated transcriptional perturbation of fibroblasts reveales an anti-fibrotic effect of IFN-γ. (A-E) Transcriptional analysis of healthy donor fibroblasts treated with IFN-γ, IL-13, or IL-17a (100ng/mL) versus untreated controls. (A) Principal component analysis (PCA) of fibroblast gene expression following cytokine treatment showing PC1, PC2, and PC3. (B) Dot plot showing overlap of cytokine-induced fibroblast gene signatures with transcriptional profiles from indicated published fibrosis datasets. (D) Enrichment of Reactome pathways across cytokine-treated fibroblasts, with the direction and strength of enrichment indicated (cytokine effect on pathway, Red = Up; Blue = Down). (D) Heatmap shows interferon-stimulated and fibrosis-associated transcripts regulated by cytokines (z-score normalized); genes overlapping with public datasets in ‘B’ are marked. Bolded genes marked with an asterisk are plotted in ‘E’. (E) Log2-transformed expression of select genes in fibroblasts treated with recombinant IFN-γ, IL-13, or IL-17a. Each point represents an independent biological replicate. Statistical significance: *p < 0.05, **p < 0.01, ns = not significant (unpaired t test or Mann-Whitney U test based on Shapiro-Wilk distribution testing). Error bars show mean ± SD.

IFN-γ selectively downregulated the expression of multiple collagen family genes, and antagonized common stem-like, wound healing, and myofibroblast genes (Fig. 6E).

Together, these data suggest that defective type 1 immunity in SSc patients may result in reduced control of fibrosis progression.

## Discussion

Analysis of lineage-defining transcription factors in individual T cells offers insights in the functional capabilities of these cells without altering their phenotype through *in vitro* restimulation. To understand differences in functional T cell subsets in peripheral blood from patients with the systemic autoimmune diseases SSc and SLE, we designed a multidimensional spectral flow cytometry panel including the lineage-defining transcription factors T-bet, GATA-3, RORγt, Bcl-6, and Foxp3, combined with phenotypical and functional receptors and additional intracellular molecules. Automated clustering analysis of the data sets generated from cohorts of lcSSc, dcSSc, and SLE patients, and healthy controls, revealed undescribed granularity in T cell subsets defined by transcription factor expression. Previously unknown features of the circulating T cell landscape emerged, distinguishing dcSSc, lcSSc, and SLE patients and healthy controls. Although SSc has been described as a disease driven by T cell-derived type 2 cytokines that exert pro-fibrotic activities on fibroblasts (20), we surprisingly did not find a clear difference in the frequencies of GATA-3-expressing T cells. Rather, we found a prominent decrease in some subsets of T-bet-expressing cells within the CD8^+^ and CD4^-^CD8^-^ γδ T cell populations, suggesting this loss may create a more permissive environment for type 2 immune cells to exert their functions in an unrestrained fashion. Decreased circulating CD8^+^ T cell numbers and dysfunctional CD8^+^ T cells have been shown in SSc (24, 51). Fuschiotti *et al.* demonstrated that CD8^+^ T cells from SSc patients contain a population of IL-13-producing cells with pro-fibrotic activity (9). Mechanistically, this was associated with cytosolic sequestering of T-bet, a known promoter of type 1 cytokine production and consequently an inhibitor of pro-fibrotic type 2 cytokines (22). Recently, Padilla *et al.* showed increased CD8^+^ tissue resident memory T cells in lungs from SSc-ILD patients, but their functional properties remain unclear (52). γδ T cells are important sources of IFN-γ and TNFα and have been proposed to play a role in autoimmune rheumatic diseases, including SSc (53).

Circulating γδ T cells numbers were shown to be lower in peripheral blood of SSc patients, but were activated and stimulated pro-collagen gene expression in fibroblasts (54), and interacted with endothelial cells (55). Moreover, γδ T cells appeared to accumulate in SSc skin (56). Our analysis of CD4^-^CD8^-^ γδ T cells provides new information about the specific subpopulations of γδ T cells that are reduced in SSc, i.e. the ones expressing T-bet (57, 58). Some authors have described γδ T cells with anti-fibrotic properties, and this may be related to their strong capacity for producing IFN-γ and TNFα, cytokines that kill aberrant fibroblasts (47, 54). The observed reductions in CD8^+^ and γδ T cells that express T-bet and are sources of IFN-γ may contribute to a tissue environment that favors type 2 immune responses and fibrosis progression.

Studies in mice further corroborate this notion since T-bet deficiency has been shown to exacerbate lung and skin fibrosis in multiple models (59–61).

Type 1 immunity in dcSSc patients may be further restrained by the increased presence of a specialized Foxp3+ Treg subpopulation expressing TIGIT. Foxp3+ Tregs expressing TIGIT have been associated with selective suppression of Th1 and Th17 responses by increasing IL-10 production in dendritic cells (15, 27), further underscoring the presence of an immune landscape characterized by compromised type 1 immune activity. Interestingly, a recent study showed that TIGIT engagement in Foxp3+ Tregs induces tissue repair functions through the secretion of the tissue growth factor amphiregulin, suggesting increased presence of these cells may contribute to excessive tissue repair leading to fibrosis (62). It is important to note here that the TIGIT-expressing Treg clusters were only increased in dcSSc patients. dcSSc and lcSSc patients differ in skin and organ involvement, as well as autoantibody profile. These distinct phenotypes are likely caused by different underlying pathophysiological mechanisms. A specific immunopathologic role of TIGIT+ Tregs in dcSSc vs lcSSc would provide a mechanism to classify dcSSc and lcSSc as disease endotypes.

Our findings also have translational implications. The overall deficiency in T cells with the capacity to produce IFN-γ in SSc patients may provide an opportunity to correct proper control of tissue repair and fibrosis by restoring specific cell subsets through cell-based therapies or biologics, an approach that may be more effective than previous attempts to treat fibrotic conditions with IFN-γ (63, 64). In mouse models, agents that promote the IFNγ/Th1 response and reduce Th2 immunity have been shown to protect against pulmonary fibrosis (65). Nevertheless, caution is warranted as T cell function is often highly dependent on the tissue cytokine microenvironment, and IFN-γ-secreting populations have been associated with both anti-fibrotic and pro-fibrotic activities in various liver diseases for example (66, 67).

For our study, we chose to extend our analysis to T cells from another systemic rheumatic disease, SLE, to reveal unique features of the SSc T cell landscape. In fact, we discovered that the severe reduction in specific type 1 T cell subsets was shared between these diseases. Nevertheless, SLE contained other, disease-specific changes such that the overall T cell landscape in SLE clearly differs from SSc. We propose that the altered balance and composition of the aberrant T cell subsets in each disease results in differential functional integration that underlies unique disease pathophysiology.

Finally, we report the surprising finding that some T cell subsets are more prone to immune-modifying treatments than others. While we did not observe significant sensitivity of T-bet or GATA-3-expressing subsets to immunosuppression, RORγt-expressing subsets appear particularly sensitive to immune-modifying therapies, and were restored to normal levels in patients treated with immunosuppressive drugs.

Whether restoration of these specific subsets contributes to symptom alleviation and slowed disease progression remains unanswered but worth exploring. Together, our study provides a map to identify T cell subsets that are the most likely impactful targets of immune modulation in SSc and SLE, revealing novel candidates for T cell-based precision therapies to repair the aberrant immune environment underlying pathology.

## Methods

### Sex as a biological variable

Our study examined both male and female tissue with similar findings reported for both sexes. The study was not powered to detect sex-based differences.

### Study participants

Blood samples were obtained from patients in BD Vacutainer® CPT™ with citrate or EDTA Blood Collection Tubes (BD Vacutainer®). All SSc patients fulfilled the 2013 revised ACR classification criteria (68). Patients were classified as having diffuse (dcSSc) or limited (lcSSc) disease according to LeRoy criteria (69). Patients and healthy controls were sex-, race- and age-matched. Diagnosis of SLE was made following EULAR/ACR criteria (70). Some patients were treated with one or more immunosuppressive drugs (Cellcept, methotrexate, tofacitinib, prednisone, and hydroxychloroquine) at the time of blood draw.

### PBMC isolation

Blood samples were collected and PBMCs isolated the same day or up to 18 hours post blood collection. PBMCs were isolated either by underlaying with Lymphoprep^TM^ (StemCell Technologies) and separating PBMCs by density dependent centrifugation, or by using a BD Vacutainer® CPT™ with citrate. PBMC samples were frozen in 50% complete RPMI 1640 medium (10% FBS, 1% non-essential amino acids, 1% HEPES, 1% PenStrep, 1% Sodium Pyruvate, 0.1% β-mercapto-ethanol) with 40% FBS and 10% DMSO, using a Mr. Frosty freezing container and placed overnight at −80°C before transferring to a liquid nitrogen tank.

### Staining for flow cytometry

For staining, frozen PBMCs were thawed in a 37°C water bath for 2 minutes. Cells were resuspended in complete RPMI 1640 medium and centrifuged at 300 x g for 10 minutes at 4°C. Supernatant was discarded and cells were re-suspended in medium and washed. Next, thawed PBMCs were washed in PBS and stained with Live/Dead stain (UV zombie, 1:800), following the manufacturer’s instructions (Biolegend). For staining of cell surface markers, cells were re-suspended in PBS + 2% FBS and Fc receptors blocked with TruStain FcX (Biolegend), followed by addition of the fluorescently-labeled antibodies specified in Suppl. Table 1A. Samples were stained for 30 minutes in the dark at 4 °C. Excess antibody was removed by washing in PBS + 2% FBS. For subsequent intracellular staining, cells were fixed in Foxp3 Fixation/Permeabilization working solution for 1 hour at room temperature in the dark. Cells were washed with permeabilization buffer, and intracellular antibody mix was added to the samples and incubated for 2 hours at room temperature in the dark. Cells were washed twice with permeabilization buffer, and resuspended in FACS buffer before being analyzed on a Cytek Aurora flow cytometer.

### T cell intracellular cytokine staining

Frozen PBMC were thawed, washed with wash buffer (RPMI + 3% FBS), resuspended in ImmunoCult™-XF T Cell Expansion Medium (StemCell Technologies, 10981) and rested overnight. The next morning, untouched T cells were isolated using pan-T cell magnetic beads (Miltenyi Biotec, 130-096-535). T cells were stimulated with phorbol 12-myristate 13-acetate (PMA) (50 ng/ml; Sigma-Aldrich, P8139) and ionomycin (1 μg/ml; Sigma-Aldrich, I06434) with brefeldin A (10 μg/ml; Sigma-Aldrich, B6542) for 5 hours. Surface staining was performed for 15 minutes at 37 °C with antibodies diluted in 0.3% BSA-HBSS (Suppl. Table 1B). For permeabilization and fixation, Cytofix/Cytoperm was used (Becton Dickinson, RUO 554714). Intracellular cytokine staining was performed for 30 minutes at 4 °C. Data were acquired on LSR Fortessa or LSRII, and analyzed using FlowJo software (Becton Dickinson, v.10.9).

### High-dimensional multiparameter full spectrum flow cytometry and algorithmic analysis

High-dimensional multiparameter FSFC was performed on the 5 laser Cytek Aurora analyzer (Cytek Biosciences). For spectral unmixing, UltraComp compensation beads (ThermoFisher) or single stained PBMCs were used, and the data analyzed with SpectroFlo (Cytek Biosciences) software, using the ordinary least square algorithm. A data processing pipeline was established using the Omiq.ai cloud computation platform. Live single cell lymphocytes were manually gated on CD3 followed by manual gating of CD4^+^, CD8^+^ and CD4^-^CD8^-^ T cells. Unmixed spectral cytometry data (FCS files) were uploaded into omiq, data were arcsinh-transformed (cofactor: 6000) and scales adjusted per signal range for each feature. All channel measurements, except CD3, CD4, CD8, PD-1, Vδ1, and Vδ2, were projected into two-dimensional space using the opt-SNE algorithm. PD-1 was not included in opt-SNE and PhenoGraph input processing because we detected substantial differences in the MFI between batches of samples that severely affected automated analysis. Nevertheless, expression of PD-1 could still be evaluated and interpreted in cluster analysis output within each experimental batch.

TCR Vδ1 and TVδ2 staining was not included in the initial batches and therefore also not in the input for automated clustering algorithm. The following parameters of the algorithm were used: random initialization, perplexity = 40, KLD minimization endpoint = 5000. For PhenoGraph, the following parameters were used: kNN = 20; distance: Euclidean; features for clustering: TIGIT, T-bet, CXCR5, CD226, RORγt, Ki-67, FOXP3, Bcl-6, GATA3, TOX, CD45RO. After the unsupervised clustering step, datapoints for each cluster were visualized as heatmap overlays over opt-SNE maps. Frequencies of individual clusters were calculated for each subject; some clusters were adjusted based on conventional expert gating. Data were plotted using Prism 8.0 (GraphPad).

### Stimulation of fibroblasts, bulk RNA-sequencing, and bioinformatic analysis

Human dermal fibroblasts from the eyelid of a single healthy donor were purchased from a commercial vendor (ZenBio; DF-F) and cultured at low passages (p = 5) using DMEM supplemented with 10% FBS, 2 mM L-glutamine, 100 U/ml penicillin, and 100 mg/ml streptomycin (Gibco™; 10938025). Fibroblasts, approximately 1-2 x 10^4^ in a 96-well flat bottom plate, were treated for 24 hours with carrier-free recombinant human IFN-γ, IL-13, or IL-17a (Biolegend; 570202, 571102, 570502) in the presence of Immunocult^TM^ CD3/CD28/CD2 Human T cell Activator (StemCell Technologies; 10970). High-quality total RNA was isolated from fibroblasts using TRIzol^TM^ Reagent (Invitrogen^TM^; 15596026). Next, cDNA was prepared using the SMART-Seq v4 Ultra Low Input RNA Kit for Sequencing (Takara). Library construction was performed using the NexteraXT DNA sample preparation kit (Illumina) using half the recommended volumes and reagents. Dual-index, single-read sequencing of pooled libraries was run on a HiSeq2500 sequencer (Illumina) with 58-base reads and a target depth of 5 million reads per sample. Base-calling and demultiplexing were performed automatically on BaseSpace (Illumina) to generate FASTQ files. The FASTQ files were processed to remove reads of zero length (fastq_trimmer v.1.0.0), remove adapter sequences (fastqmcf tool v.1.1.2), and perform quality trimming from both ends until a minimum base quality ≥ 30 (FASTQ quality trimmer tool v.1.0.0). Reads were aligned to the human reference genome (build hg38) with TopHat (v.1.4.0) and read counts per Ensembl gene ID were quantified with htseq-count (v.0.4.1) (71, 72). Quality metrics for the FASTQ and BAM/SAM files were generated with FastQC (v.0.11.3) and Picard (v.1.128). Processing of FASTQ and BAM/SAM files was executed on the Galaxy workflow platform of Globus genomics. Statistical analysis of gene expression was assessed in the R environment (v.4.4.2). All samples passed a set of pre-established quality control criteria: total number of fastq reads > 1 x 10^6^; mapped reads > 70%; median CV coverage < 0.85. Randomization was not applicable as no treatment or intervention groups were included in the study. Blinding was not applicable as no treatment groups were compared.

### Additional bioinformatic analyses

Differential expression and pathway enrichment analyses were performed in R. To assess differential expression, the limma package (v.3.58.1) was used (73). The limma voomWithQuality weights transformation of TMM normalized counts was used to fit a linear model of gene expression as a function of recombinant cytokine treatment followed by empirical bayes smoothing of standard errors. A log2 expression fold change of at least 1 in magnitude and an FDR of less than 0.05 were used as cutoffs to define differentially expressed genes.

Overlap of differentially expressed genes and published fibrosis signatures was assessed using a hypergeometric test. Pathway enrichment analysis was performed using the enrichR package (v.3.2) (74) with the Reactome 2022 database (75) to query pathways associated with cytokine-induced genes.

### Statistics

Statistical analyses were performed using Prism Version 9.2.0 (GraphPad). Specific statistical analyses that were used for data in each figure are noted in the legends, i.e. Kruskal-Wallis test with Dunn’s test for multiple comparisons, one-way analysis of variance (ANOVA) followed by post hoc Tukey’s test for multiple comparisons, two-way ANOVA followed by post hoc Tukey’s test for multiple comparisons and Student’s t-test. Statistical significance was determined based on p ≤0.05. For all figures, data are shown as mean ± standard error of the mean (SEM). Study approval All samples and clinical data were collected upon written informed consent under a protocol approved by the Institutional review board (IRB) at Boston University.

Additional samples were obtained upon written informed consent at Virginia Mason Franciscan Health (Seattle, WA, USA), according to protocols approved by the Institutional Review Board of Benaroya Research Institute (Seattle, WA, USA)(48).

## Data availability

All datasets generated and analyzed in this study are available from the corresponding authors on reasonable request. All gene expression data sets generated in this study are available to the public in the Gene Expression Omnibus, GSE305982. Computer code generated and used for analysis in this study is available in Github: https://github.com/BenaroyaResearch/VolfsonAlteredTCellLandscapeInSScAndSLE

## Author contributions

All authors contributed significantly to the study. VVS conducted experiments, analyzed data and contributed to writing the manuscript; HAD performed bioinformatics analysis; MLF conducted experiments; IRR collected and provided patient samples and data; AMB collected and provided patient samples and data; DJC designed research studies and supervised the study; PAM conducted experiments, analyzed data, supervised the study, and contributed to writing the manuscript; ACB developed data analysis methods, analyzed data and supervised the study; HD conceptualized and supervised the study, and wrote the manuscript.

## Acknowledgements

We are grateful to Eric Stratton and Drs. Robert Lafyatis, Marcin Trojanowski and Hanni Menn-Josephy, and members of the Lupus Multidisciplinary Program at Boston Medical Center for assistance with human subject recruitment and Internal Review Board matters at Boston University Medical Campus. We thank Dr. Maria Trojanowska for general support and expertise provided for this study. We acknowledge the technical support from Dr. Jennifer Snyder-Cappione and the Boston University Flow Cytometry Core Facility.

Furthermore, we are grateful to Kassidy Benoscek-Narag, Sylvia Posso, Thien-Son Nguyen, and the Clinical Research Center coordinators in the Benaroya Research Institute (BRI) Center for Interventional Immunology for human subject recruitment, sample collection and screening through the BRI Biorepository (RRID:SCR_026967); the BRI Cell and Tissue Analysis (RRID:SCR_026327) and Genomics (RRID:SCR_026658) facilities for technical assistance and expertise. BRI cytometry and genomics equipment used in this study were generously supported by the M.J. Murdock Charitable Trust. We are very grateful to the patients at both sites for their participation in this study.

