## Supplemental Data for "The altered T cell landscape in Systemic Sclerosis patients is characterized by dysfunctional type 1 immunity"

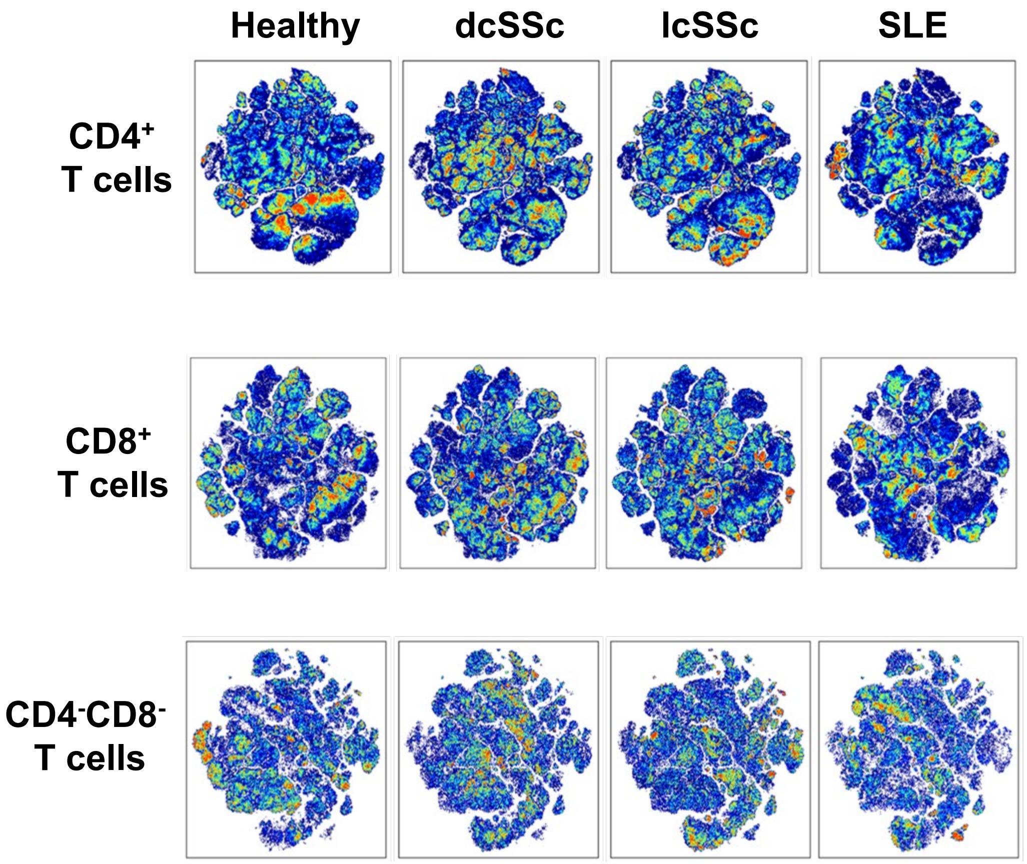


**Suppl. Figure 1: Cell distribution comparison between healthy subjects, and dcSSc, lcSSc and SLE patients**

Opt-SNE maps of CD4+, CD8+ and $CD4^{-}CD8^{-}$ T cells overlaid with cell densities for healthy controls, dcSSc, lcSSc and SLE patients. Each cohort was normalized to contain the same number of cells per cohort for each T cell lineage: $1x10^{6}$ cells for CD4+ and CD8+ T cells, and $2.5 \times 10^{5}$ for $CD4^{-}CD8^{-}$ T cells. Data represents all samples concatenated.

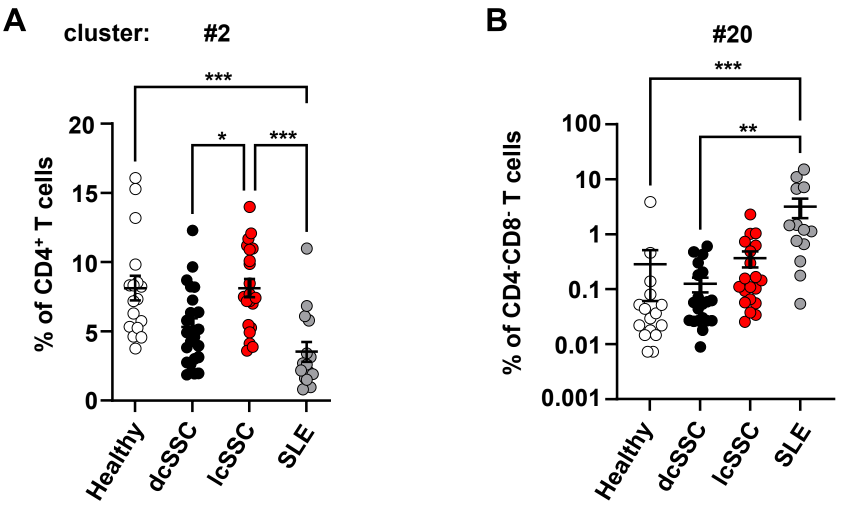


**Suppl. Figure 2: Aberrant features of circulating Tfh and Bcl-6$^{+}$ T cells in SSc and SLE**

Frequencies of (A) Tfh and (B) Bcl-6+ clusters within CD4+ (cluster 2) and CD4-CD8- (cluster 20) parental populations of healthy controls (n=17), dcSSc (n=23), lcSSc (n=21), SLE (n=15) patients as shown in Fig. 1D, F. Bar graphs show percentages of gated cells of indicated clusters with mean and SEM, calculated as in Fig. 2. Each symbol represents a blood sample from an individual donor. Data were statistically analyzed with the Kruskall-Wallis test followed by Dunn's multiple comparisons test. *p<0.05, **p<0.01, ***p<0.001

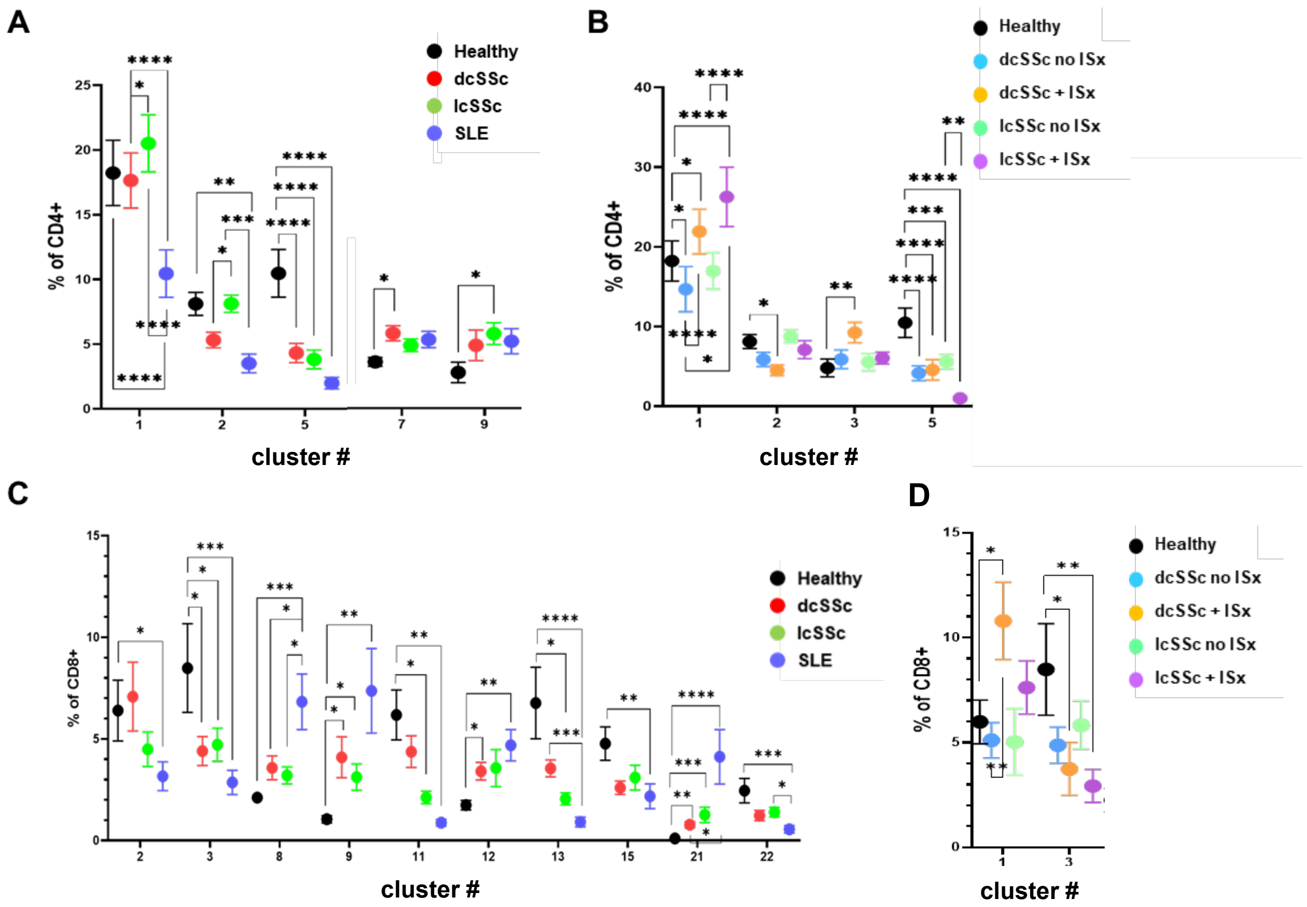


**Suppl. Figure 3: Additional differences in frequencies of clusters not defined by transcription factor expression in peripheral blood**

(A) Comparison of frequencies of selected clusters within the CD4+ T cell compartment identified in Fig. 1D and analyzed as in Fig. 2. Data is shown as the mean of healthy controls (n=17), dcSSc (n=23 ), lcSSc (n=21), and SLE (n=15) patients +/- SEM. (B) Impact of immunosuppressant treatment (ISx) on frequencies of selected clusters within CD4+ T cells. (C) Comparison of frequencies of selected clusters within the CD8+ T cell compartment identified in Fig. 1E and analyzed as in Fig. 2. (D) Impact of immunosuppressant treatment (ISx) on frequencies of selected clusters within CD8+ T cells. Data were statistically analyzed with two-way ANOVA test. *$p<0.05$, **$p<0.01$, ***$p<0.001$, ****$p<0.0001$

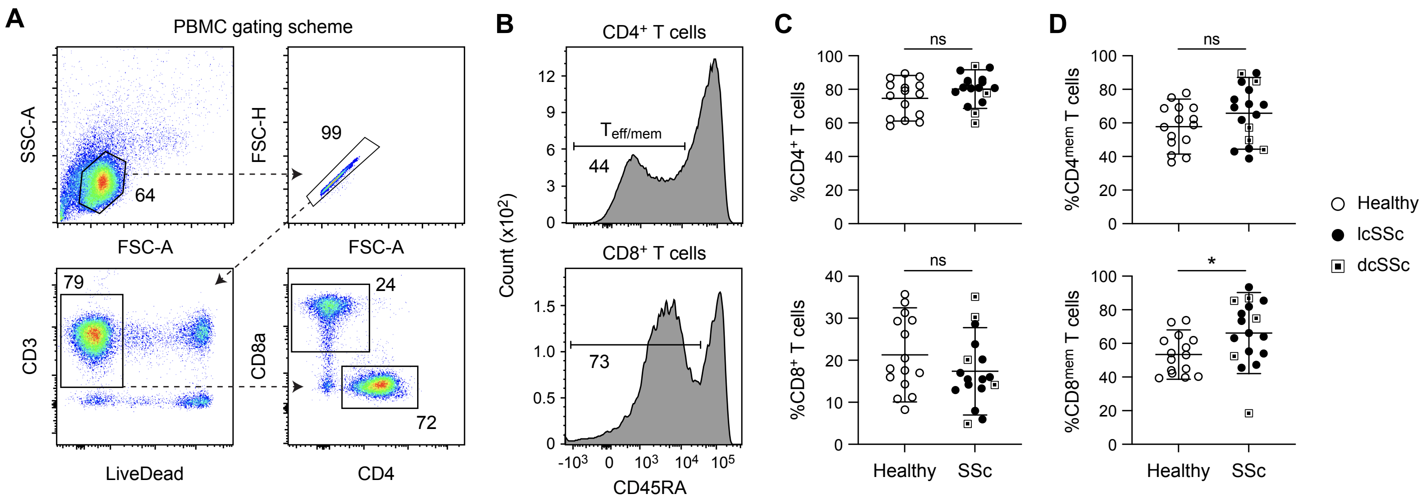


**Suppl. Figure 4: Gating strategy for peripheral blood T cells from SSc subjects.** (A) Flow cytometry gating for identification of CD3+CD4+ and CD3+CD8+ T cells from PBMCs, including singlet, viability, and lineage gating steps. (B) Representative CD45RA histograms for CD4+ and CD8+ T cells showing memory (CD45RAlo) versus naïve populations. (C-D) Frequencies of peripheral T cell subsets in healthy donors and SSc patients. (C) Proportion of CD4+ and CD8+ T cells among CD3+ T cells. (D) Percentage of CD45RAlo memory T cells among CD4+ (top) and CD8+ (bottom) subsets.

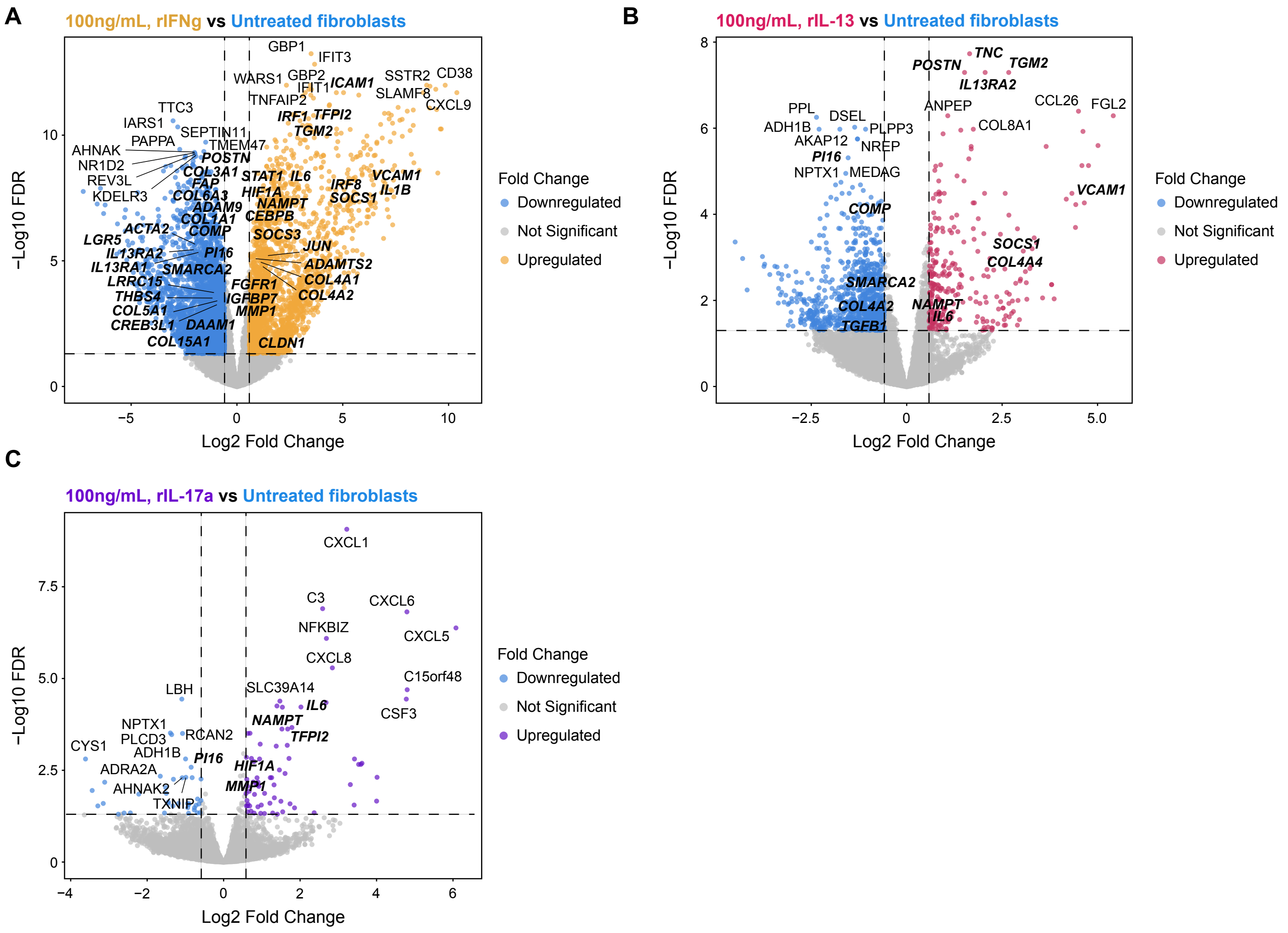


**Suppl. Figure 5**: **Cytokine-specific gene regulation in dermal fibroblasts.** (A-C) Volcano plots showing differentially expressed genes in fibroblasts treated for 24 hours with 100ng/mL recombinant IFN-γ (A), IL-13 (B), or IL-17a (C), compared to untreated fibroblasts. The top 10 genes up- or down-regulated per condition ranked by -log10 FDR are shown. Bold italicized genes are differentially regulated and overlap with the heatmap in Figure 6D. Statistical significance: *p < 0.05, **p < 0.01, ns = not significant (unpaired t test or Mann-Whitney U test based on Shapiro-Wilk distribution testing).

**A**

| Specificity | Fluorophore | Vendor | Catalog # | Clone | Staining concentration |
|---|---|---|---|---|---|
| L/D | UV Zombie | Biolegend | 423107 | - | 1:800 |
| CD25 | BUV395 | BD | 564034 | 2A3 | 1:50 |
| CD127 | BUV737 | BD Biosciences | 612795 | HIL-7R-M21 | 1:50 |
| PD-1 | BV421 | Biolegend | 329920 | EH12.2H7 | 1:50 |
| CD3 | BV510 | Biolegend | 317332 | OKT3 | 1:100 |
| TIGIT | BV605 | Biolegend | 372712 | A15153G | 1:50 |
| T-bet | BV711 | Biolegend | 644820 | 4B10 | 1:50 |
| CXCR5 | BV750 | Biolegend | 356942 | J252D4 | 1:25 |
| CD226 | BV786 | BD Biosciences | 742497 | DX11 | 1:50 |
| RORγt | AF488 | BD Biosciences | 563621 | Q21-559 | 1:20 |
| Ki-67 | AF532 | eBioscience/ ThermoFisher | 58-569-882 | SolA15 | 1:200 |
| CD4 | PerCP-e710 | eBioscience/ ThermoFisher | 50-112-9403 | SK3 | 1:3200 |
| FOXP3 | PE | eBioscience/ ThermoFisher | 12-4776-42 | PCH101 | 1:100 |
| Bcl-6 | PE/Dazzle594 | Biolegend | 358510 | 7D1 | 1:40 |
| GATA3 | PE-Cy7 | eBioscience/ ThermoFisher | 25-9966-42 | TWAJ | 1:5 |
| TOX | APC | Miltenyi | 130-118-335 | REA473 | 1:100 |
| CD45RO | Alexa700 | BD Biosciences | 561136 | UCHL1 | 1:50 |
| CD8 | APC-Fire/810 | Biolegend | 344764 | SK1 | 1:100 |
| Vδ1 | APC-Vio770 | Miltenyi | 130-120-438 | REA173 | 1:100 |
| Vδ2 | PerCP | Biolegend | 331410 | B6 | 1:50 |

**B**

| Specificity | Fluorophore | Clone | Vendor | Catalog # | RRID # | Staining concentration (ug/mL) |
|---|---|---|---|---|---|---|
| Identification and cytokine staining of blood-derived T cells (Figure 5, Supplemental Figure 4) | - | - | - | - | - | - |
| CD3 | BUV395 | OKT3 | Thermo Fisher Scientific | 363-0037-42 | RRID:AB_2925248 | 1 |
| GM-CSF | BV421 | BVD2-2C11 | BD Biosciences | 562930 | RRID:AB_2737899 | 1 |
| CD8a | BV510 | RPA-T8 | Biolegend | 301048 | RRID:AB_2561942 | 1 |
| CD45RA | BV605 | HI100 | Biolegend | 304133 | RRID:AB_11126164 | 0.625 |
| CD4 | BV711 | OKT4 | Biolegend | 317439 | RRID:AB_11219404 | 1 |
| CLA | FITC | HECA-452 | Biolegend | 321306 | RRID:AB_492898 | 4 |
| IL-13 | PE | JES10-5A2 | Biolegend | 501903 | RRID:AB_315198 | 0.83 |
| IL-22 | PE/Cy7 | 22URTI | Thermo Fisher Scientific | 25-7229-42 | RRID:AB_10853346 | 0.5 |
| CD103 | APC | Ber-ACT8 | Biolegend | 350216 | RRID:AB_2563907 | 2.5 |
| IFNg | AlexaFlour700 | 4S.B3 | Biolegend | 502519 | RRID:AB_528920 | 5 |
| Fixable LiveDead | NearIR | n/a | Thermo Fisher Scientific | 65-0865-14 | - | 1:1000 dilution |

**Supplementary Table 1:** Composition of (A) 20-color multiparameter spectral cytometry panel for immunophenotyping, and (B) antibody panel used for intracellular cytokine staining.
